# *In vitro* synergistic action of TAT-RasGAP_317-326_ peptide with antibiotics against Gram-negative pathogens

**DOI:** 10.1101/2022.09.26.509445

**Authors:** Grazia Vizzarro, Nicolas Jacquier

## Abstract

**Objectives:** Multidrug-resistant (MDR) bacteria are a continuously increasing threat for medicine, causing infections recalcitrant to antibiotics. Antimicrobial peptides (AMPs) were identified as alternatives to antibiotics, being naturally occurring short peptides and part of the innate immune system of a vast majority of organisms. However, the clinical application of AMPs is limited by suboptimal pharmacokinetic properties and relatively high toxicity. Combinatorial treatments using AMPs and classical antibiotics may decrease the concentrations of AMPs required for bacterial eradication, thus lowering the side effects of these peptides.

**Methods:** Here, we investigate the *in vitro* efficiency of combinations of the recently described antimicrobial peptide TAT-RasGAP_317-326_ with a panel of commonly used antimicrobial agents against three Gram-negative bacteria: *Escherichia coli, Pseudomonas aeruginosa* and *Acinetobacter baumannii* using checkerboard and time-kill assays.

**Results:** We identified synergistic combinations towards all three bacteria and demonstrated that these combinations had an increased bactericidal effect compared to individual drugs. Moreover, combinations were also effective against clinical isolates of *A. baumannii*. Finally, combination of TAT-RasGAP_317-326_ and meropenem had a promising antibiofilm effect towards *A. baumannii*.

**Conclusion:** Taken together, our results indicate that combinations of TAT-RasGAP_317-326_ with commonly-used antimicrobial agents may lead to the development of new treatment protocols against infections caused by MDR bacteria.

## 1. Introduction

Infections caused by Gram-negative multidrug-resistant (MDR) bacteria are an increasing concern for modern medicine. *Pseudomonas aeruginosa* and *Acinetobacter baumannii* have been classified by WHO and NHS as especially concerning for human health. *P. aeruginosa* is one of the primary causes of nosocomial and ventilator-associated pneumonia, with a high mortality rate in immunocompromised, transplanted, or cystic fibrosis patients [1]. Antibiotic treatment of *P. aeruginosa* infections is difficult due to its low outer membrane permeability and the presence of a wide variety of both intrinsic and acquired resistance mechanisms [2, 3]. Likewise, *A. baumannii* is an increasingly concerning human pathogen, causing hospital- and community-acquired infections, including pneumonia, bacteremia, endocarditis, skin and urinary tract infections, and meningitis [4]. These infections are common in immunocompromised, ventilated and intensive care unit patients. Carbapenems are commonly used to treat *A. baumannii* infections, and carbapenem-resistant *A. baumannii* have emerged rapidly and have been detected worldwide [5-7]. This rapid development of resistance may be linked to the high genome plasticity of this bacterial species [8-10]. *A. baumannii* is also capable of surviving in healthcare environments, despite routine disinfection regimes, as it is resistant to desiccation and oxidative stress [11].

Both *P. aeruginosa* and *A. baumannii* can develop drug-tolerant communities called biofilms on abiotic surfaces and tissues [12]. These biofilms contain bacteria encased in a matrix composed of extracellular polymeric substances, including polysaccharides, extracellular DNA and proteins, among others. Biofilm formation results in antimicrobial treatment failure, as bacteria present within a biofilm show strongly increased tolerance to antibiotics compared to planktonic bacteria (free-living bacterial cells) [13]. Novel antimicrobials and new treatment strategies are thus required to efficiently fight infections caused by drug-resistant and biofilm-forming bacteria.

Antimicrobial peptides (AMPs) are promising antibacterial agents. Several of them are effective against MDR bacteria [14, 15] and possess antibiofilm activity [16]. Different factors have hindered the progress of AMPs in preclinical or clinical trials against MDR bacteria : suboptimal pharmacokinetic properties and relatively high toxicity to mammalian cells, among others [17]. We recently reported the antibacterial and antibiofilm potential of the TAT-RasGAP_317-326_ peptide against a range of pathogenic bacteria, including *P. aeruginosa, A. baumannii* and *Escherichia coli* [18, 19]. TAT-RasGAP_317-326_ minimal inhibitory concentrations (MICs) vary according to the bacterial species (8-16 μg/mL for *A. baumannii*, 16-32 μg/mL for *E. coli*, and 64-128 μg/mL for *P. aeruginosa*) and to environmental factors such as medium composition [20]. Moreover, clinical application of this peptide is hindered by suboptimal biodistribution and rapid clearance [18, 21]. It is thus necessary to develop strategies to increase the efficiency of this peptide in order to overcome limitations caused by its pharmacokinetic properties.

In this study, we attempted at combining TAT-RasGAP_317-326_ with a selection of nine antimicrobial agents. This allowed to identify candidate drugs, polymyxin B, aztreonam and gentamicin, showing a promising interaction with TAT-RasGAP_317-326_ towards *E. coli, P. aeruginosa* and *A. baumannii*, respectively. We could then show using checkerboard assays that these drugs had a synergistic interaction with TAT-RasGAP_317-326_. We then investigated the effect of combinations on *A. baumannii* biofilm and could determine that combining TAT-RasGAP_317-326_ with meropenem was efficient at both inhibiting biofilm formation and affecting viability of established biofilm.

## 2. Methods

### 2.1 Bacterial strain and growth conditions

The bacterial strains used are described in Supplementary Table S1. All the strains were cultured in LB medium (10 g/L tryptone, 5g/L yeast extract, 10g/L NaCl) at 37°C and 200 rpm shaking and on LB agar plates at 37 °C statically.

### 2.2 Peptides and antibiotics

The retro-inverso all-D peptide TAT-RasGAP_317-326_ (amino acid sequence DTRLNTVWMWGGRRRQRRKKRG) was synthesized by SBS Genetech (Beijing, China) and stored at -20°C. Polymyxin B sulfate salt, ceftriaxone disodium salt, erythromycin, aztreonam and meropenem were purchased from Sigma-Aldrich (Burlington, MA), ampicillin sodium salt, ciprofloxacin, gentamicin, and tetracycline from Applichem (Darmstadt, Germany). All antibiotics were resuspended in water and stored at -20°C, with the exception of azithromycin and tetracyclin, which were resuspended in ethanol and stored at -20°C and of aztreonam, which was resuspended in DMSO and stored at -80°C.

### 2.3 Minimal inhibitory concentration (MIC) measurement

The antimicrobial activity of compounds was measured using the Microtiter Broth Dilution method performed on 96-well plates. Bacteria were pre-cultured in LB overnight and then diluted to an OD600 of 0.1 and grown at 37°C with shaking for 1 hour to allow re-growth to log-phase. Then, the bacterial suspension was diluted 100 times in LB medium. The antimicrobial agents were serially two-fold diluted in LB medium in a 96-well plate in a final volume of 100 µL, and 10 µL of bacterial suspension was added into each well. The plate was incubated at 37°C for approximately 16 hours. Bacterial growth was quantified by OD590 measurement on a FLUOstar Omega Microplate Reader (BMG Labtech, Ortenberg, Germany). The MIC value was defined as the lowest concentration of an antimicrobial agent required to inhibit 90% of microbial growth. The IC50 value indicates the concentration that inhibits the growth by 50% in comparison to the untreated control and was determined by interpolation fitting a sigmoid curve to the results on GraphPad Prism 9.1.0.

### 2.4 Checkerboard assay

Synergism or additivity between antimicrobial agents were investigated in 12 × 12 checkerboard assays. Bacteria were prepared as described for MIC measurement. Two-fold serial dilutions of tested antimicrobial drugs and TAT-RasGAP_317-326_ were performed respectively across the rows and columns of the 96-well plate in a final volume of 100 µL of LB medium, followed by the addition of 10 µL of prepared bacterial inoculum. The plate was then incubated statically at 37°C for approximately 16 hours, OD590 was measured and MICs were determined. Synergistic, additive or antagonistic interactions of the antimicrobial agents were determined by calculating a fractional inhibitory concentration index (FICI) [22] with the following formula:

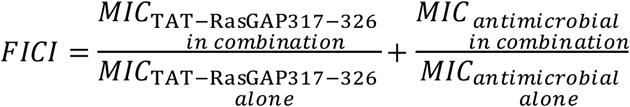

The combination with the lowest FICI value was considered as optimal (FICIopt). FICIopt ≤ 0.5 indicates a synergism, 0.5 < FICIopt ≤ 4 additivity or indifference and FICIopt > 4 antagonism. Each experiment was performed in triplicate. In case of discrepancy between the three replicates, interaction was attributed to a category when at least two of the three replicates gave FICI values consistent with this category.

### 2.6 Colony forming unit (CFU) measurements

The bactericidal activity of the selected combinations was evaluated by Colony forming unit (CFU) measurements after addition of the corresponding antimicrobial combination. Overnight bacterial cultures were diluted to 0.1 OD600 and grown for 1 hour with 200 rpm shaking at 37°C. Next, antimicrobial agents were added alone or in combination to 500 µL of bacterial suspension. 10 µL of viable bacteria were retrieved 2 or 6 hours after drug addition, and serially diluted 10-fold. 10 µL per dilution were plated on LB agar plate and grown at 37°C overnight. Colonies were counted, and CFU calculated.

### 2.7 Biofilm formation inhibition assay

Biofilm inhibition assay was performed as described earlier [19]. Briefly, bacteria were grown overnight in 5 mL of LB, diluted 1:50 in fresh media and grown for an additional 3-4 hours. Bacterial cultures were washed twice with phosphate-buffered saline (PBS) followed by centrifugation at 4000 x *g* for 10 minutes and then adjusted to 0.2 OD600 in BM2 medium (62 mM potassium phosphate buffer, 7 mM ammonium sulfate, 10 μM iron sulfate, 0.4% (w/v) glucose, 0.5% (w/v) casaminoacids, 2 mM magnesium sulfate) [23]. The antimicrobial agents alone or in combination, together with bacteria (0.1 OD600) were added in a polypropylene 96-well plate (Greiner, Frickenhausen, Germany) in a final volume of 100 µL. Biofilms were allowed to form for 24h at room temperature.

To estimate the biofilm biomass, crystal violet (Sigma) staining was used. Biofilms were washed with deionized water and stained for 10 minutes with 125 µL of crystal violet solution diluted to 0.1%. After washing twice with deionized water, 125 μL of 30% acetic acid was added and incubated for 10 minutes at room temperature. 125 μL of solution was then transferred to a new 96-well plate and the absorbance was measured at 590 nm using a FLUOstar Omega Microplate Reader.

In parallel, bacterial metabolism was measured using resazurin fluorescence to estimate bacterial viability, as described earlier [24]. Biofilm was washed twice with PBS and incubated for 90 minutes with 4 μg/mL resazurin (Sigma). Fluorescence was measured with a FLUOstar Omega Microplate Reader with Ex/Em = 540/580 and gain = 500. Combinations of drugs were tested similarly as described for checkerboard assays. The MBIC90 (minimal biofilm inhibitory concentration) was defined as the lowest concentration of the antimicrobial agent alone or in combination that reduced the formation of biofilm (viability or biomass) by ≥ 90%. The biofilm fractional inhibitory concentration index (BFICI) has been determined similarly as the FICI described in Checkerboard assay section and similar thresholds for synergism/additivity/antagonism were applied.

### 2.7 Established biofilm eradication treatment

Cells were cultured as described for biofilm formation inhibition and then adjusted to 0.1 OD600 in BM2 medium. 100 µl of bacterial solution were added to each well of a polypropylene 96-well plate, and biofilms were allowed to form for 24 hours. The established biofilms were then washed with PBS and incubated with 100 µL of the antimicrobial agent or the combination in BM2 medium for 24 hours at room temperature. The bacterial viability and the biofilm biomass were measured as described above, and the MBEC90 (minimal biofilm eradication concentration) was defined as the concentration of the antimicrobial agents alone or in combination that reduces the biofilm viability or biomass by ≥ 90%.

## 3. Results

### 3.1 Preliminary screening identifies drugs positively interacting with TAT-RasGAP_317-326_ peptide on laboratory strains of *E. coli, P. aeruginosa* and *A. baumannii*

We assembled a library of 9 antimicrobial drugs: azithromycin, aztreonam, ceftriaxone, ciprofloxacin, erythromycin, gentamicin, meropenem, tetracycline and polymyxin B, covering the main families of commonly used antimicrobial agents (Supplementary Table S2) and exposed laboratory strains *E. coli* MG1655, *P. aeruginosa* PA14 and *A. baumannii* ATCC 19606 to serial dilutions of the TAT-RasGAP_317-326_ peptide in combination with a concentration of these drugs corresponding approximately to the half of the minimal inhibitory concentration (MIC)(Supplementary Table S1 and Fig. S1). We then determined for each bacterial species which antimicrobial agent had the most influence on TAT-RasGAP_317-326_ MIC (Fig 1). We thus selected polymyxin B for *E. coli*, aztreonam for *P. aruginosa* and gentamicin for *A. baumanni* for further studies. Additionally, meropenem, which is an important treatment against infections caused by *A. baumannii* was also selected.

**Figure 1:**
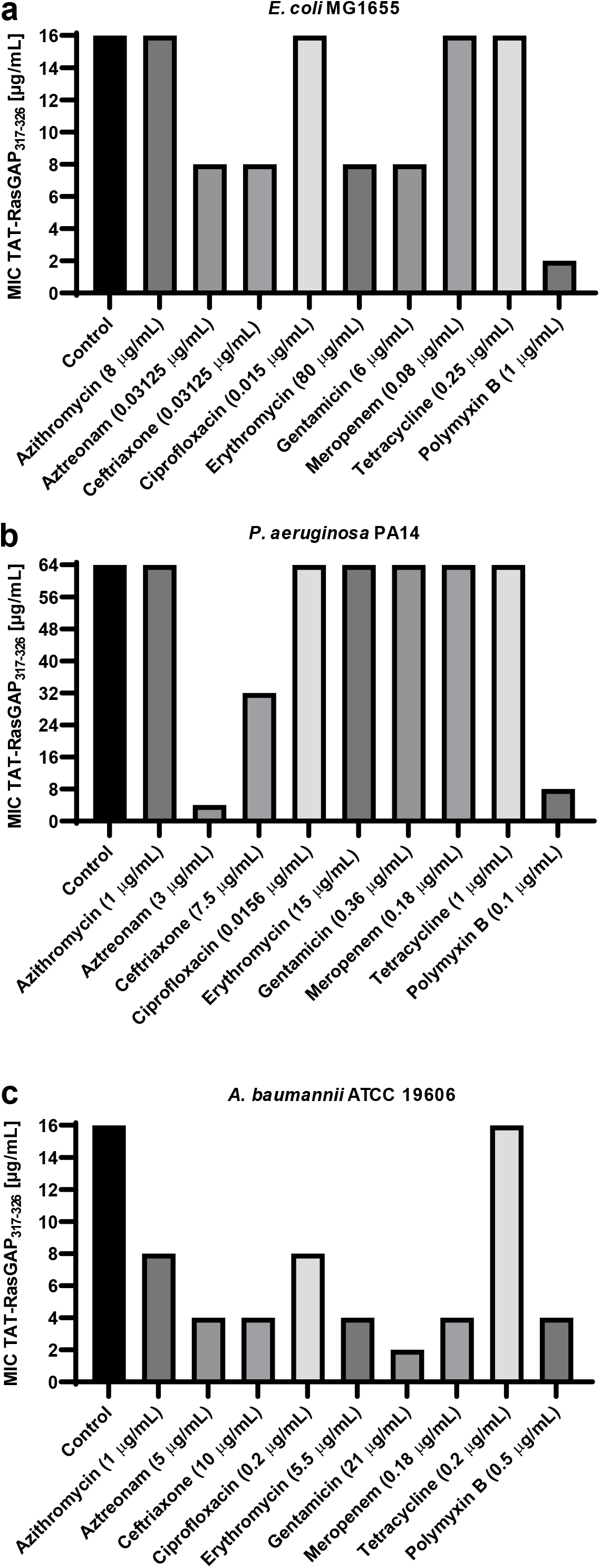
Screening of combinations between TAT-RasGAP_317-326_ and a selection of antimicrobial agents. Measurements of minimal inhibitory concentration (MIC) of TAT-RasGAP_317-326_ were performed in presence of the indicated antimicrobial agents, with concentrations corresponding approximately to the IC50 of each drug. The indicated bacteria,*E. coli* MG1655 **(a)**, *P. aeruginosa* PA14 **(b)** and *A. baumannii* ATCC 19606 **(c)** were grown in 96 well plates in presence of serial dilutions of TAT-RasGAP_317-326_ for 16 hours and OD590 was measured. MIC was determined as the minimal concentration causing inhibition of visible growth.

### 3.2 Polymyxin B, aztreonam and gentamicin in combination with TAT-RasGAP_317-326_ show a synergistic effect against *E. coli, P. aeruginosa and A. baumannii*, respectively

Synergistic interaction’s assessment of the four combinations selected during our initial screening was achieved by performing checkerboard assays and calculating the Fractional Inhibitory Concentration Index (FICI) for each pair of compounds. Classical thresholds were applied: FICI ≤ 0.5 for synergism, FICI > 0.5 and ≤ 4 for additivity and FICI > 4 for antagonism [25]. The combination of TAT-RasGAP_317-326_ and polymyxin B showed a FICI ≤ 0.5 toward both *E. coli* MG1655 and ATCC25922 (Table 1 and Supplementary Fig. S2). Similar FICI were also measured for the combination of TAT-RasGAP_317-326_ and aztreonam against *P. aeruginosa* strains PA14 and PA01 (Table 1 and Supplementary Fig. S3) and the combination of TAT-RasGAP_317-326_ and gentamicin against *A. baumannii* ATCC19606 (Table 1 and Supplementary Fig. S4a). The combination of TAT-RasGAP_317-326_ and meropenem did not show a synergistic interaction, exhibiting additivity instead (0.5 < FICI ≤ 0.75, Table 1 and Supplementary Fig. S4b).

**Table 1:** Checkerboard assays performed on laboratory strains of *E. coli, P. aeruginosa* and *A. baumannii* show that the combinations tested are synergistic (FICI ≤ 0.5: TAT-RasGAP_317-326_ and polymyxin B on *E. coli*, TAT-RasGAP_317-326_ and aztreonam on *P. aeruginosa* and TAT-RasGAP_317-326_ and gentamicin on *A. baumannii*) or additive (0.5 ≤ FICI ≤ 1: TAT-RasGAP_317-326_ and meropenem on A. baumannii). Three independent replicates were performed for each combination and MICs of the individual antimicrobial agents alone and in combination are presented. Complete checkerboard results are shown in Supplementary Figures S2, S3 and S4. Cells containing two values indicate that all three replicates gave a value that falls within the indicated range.

| Strain | MIC Alone (µg/mL) |  | MIC in Combination (µg/mL) |  | FICI |  |  |
| --- | --- | --- | --- | --- | --- | --- | --- |
|  | TAT-RasGAP<sub>317-326</sub> | Polymyxin B | TAT-RasGAP<sub>317-326</sub> | Polymyxin B |  |  |  |
| *E. coli* MG1655 | 16-32 | 2-4 | 4-8 | 0.5-1 | 0.375 | 0.5 | 0.5 |
| *E. coli* ATCC25922 | 64 | 2-8 | 16 | 0.125-1 | 0.3125 | 0.375 | 0.375 |
|  | TAT-RasGAP<sub>317-326</sub> | Aztreonam | TAT-RasGAP<sub>317-326</sub> | Aztreonam |  |  |  |
| *P. aeruginosa* PA14 | 64-128 | 8 | 16-32 | 0.5-2 | 0.375 | 0.375 | 0.5 |
| *P. aeruginosa* PA01 | 128 | 2-4 | 16-32 | 0.5 | 0.375 | 0.375 | 0.5 |
|  | TAT-RasGAP<sub>317-326</sub> | Gentamicin | TAT-RasGAP<sub>317-326</sub> | Gentamicin |  |  |  |
| *A. baumannii* ATCC 19606 | 16 | 64 | 2-4 | 8-16 | 0.375 | 0.375 | 0.5 |
|  | TAT-RasGAP<sub>317-326</sub> | Meropenem | TAT-RasGAP<sub>317-326</sub> | Meropenem |  |  |  |
| *A. baumannii* ATCC 19606 | 16-32 | 0.25-0.5 | 4-8 | 0.125-0.25 | 0.5 | 0.75 | 0.75 |

### 3.3 Synergistic combinations have a significant impact on bacterial viability

Checkerboard assays measure the synergistic effect of drug combinations on bacterial growth since turbidity of the medium is measured. We wanted to go a step further and determine if the synergistic combinations we identified could efficiently kill bacteria using Colony Forming Unit (CFU) measurements. First, we measured the minimal biocidal concentrations of individual drugs required to kill 99.9 % of bacteria after 2 hours of incubation (MBC99.9). This corresponds to a 10-3 fold change of CFU as depicted in Supplementary Figure S5. Interestingly, even high concentrations of TAT-RasGAP_317-326_ and aztreonam were unable to kill the 99.9 % of *P. aeruginosa* PA14. Similarly, meropenem only had a limited effect on *A. baumannii* ATCC 19606 viability in our settings. However, an increased killing effect was observed when drugs were combined. Specifically, the combination of TAT-RasGAP_317-326_ and polymyxin B on *E. coli* (24 μg/mL and 2 μg/mL respectively, Fig. 2a) was able to kill more than 99.9 % of bacteria. An enhanced effect was also observed upon combination of TAT-RasGAP_317-326_ and aztreonam against *P. aeruginosa* (32 μg/mL and 2 μg/mL respectively, Fig. 2b). Regarding *A. baumannii*, combination of TAT-RasGAP_317-326_ with gentamicin (8 μg/mL and 32 μg/mL respectively), but not with meropenem, was able to efficiently kill more than 99.9 % of bacteria after 2 hours of incubation (Fig. 2c and 3a). We hypothesized that the lack of effect of combination with meropenem might be due to the slow action of meropenem. We thus incubated bacteria with combination of TAT-RasGAP_317-326_ and meropenem for 4 and 6 hours and observed a significant increased bactericidal efficiency of the combination on *A. baumannii* in function of the time of incubation with the antibiotics (Fig. 3a).

**Figure 2:**
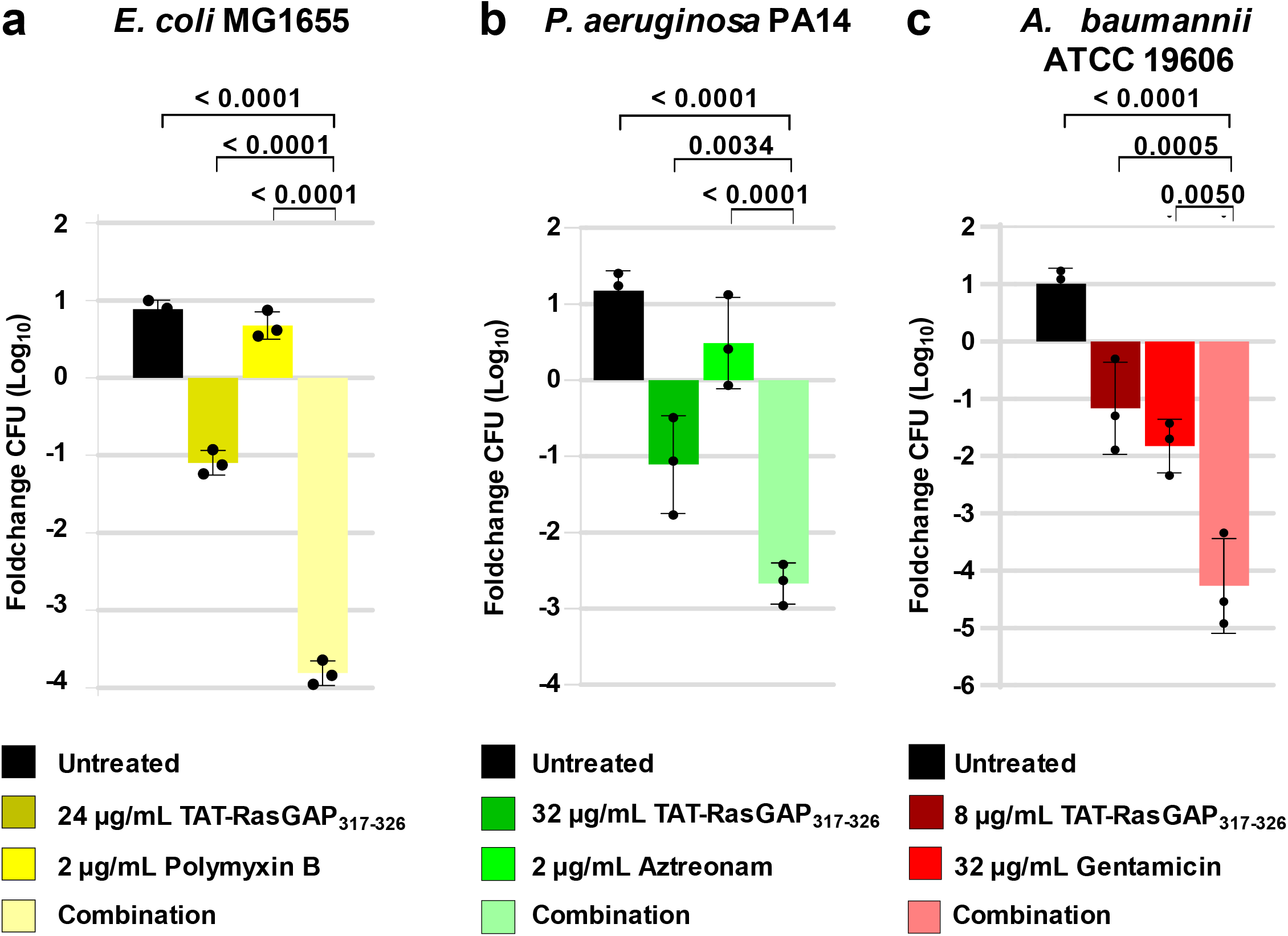
Combinations of two compounds have an increased bactericidal activity compared to individual drugs but do not result in the complete killing of bacteria. Overnight cultures of *E. coli* MG1655 **(a)**, *P. aeruginosa* PA14 **(b)** and *A. baumannii* ATCC19606 **(c)** were diluted to 0.1 OD600 and grown for one hour at 37°C before the addition of antimicrobial agents alone or in combinations, as indicated. Incubation was continued and samples were taken at 0h and 2h after addition of the drugs. Serial dilutions of these samples were plated on LB agar plates and number of Colony Forming Units per mL (CFU/mL) was quantified. Fold change between CFU/mL at each time point and the initial value (time 0) was calculated and is shown as Log10 values. Error bars indicate standard deviation of triplicates. P values were calculated using two-way Anova in Graphpad Prism.

**Figure 3:**
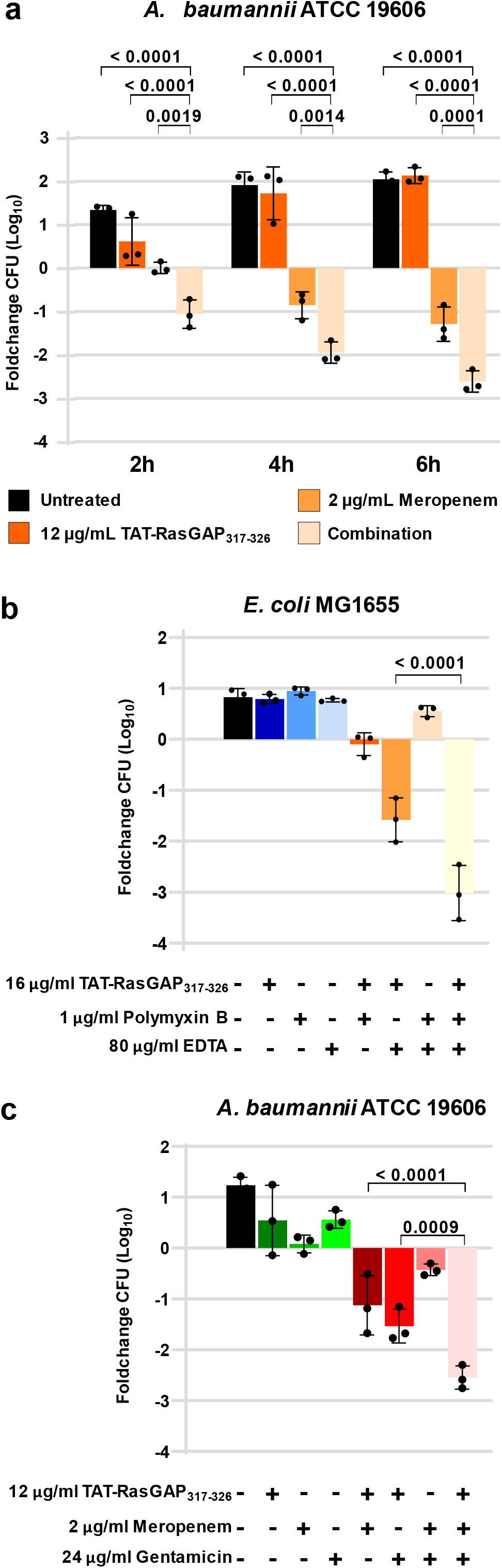
Changes in time of incubation and three-component combinations have increased killing efficiency. **(a)** Experiment was performed as for Figure 2c, with the indicated drugs and times of incubation. **(b)** Experiment was performed as for Figure 2a, in presence (+) or absence (-) of the indicated concentrations of TAT-RasGAP_317-326_, polymyxin B and of the Calcium chelator EDTA. **(c)** Experiment was performed as in Figure 2c with combinations of the indicated concentrations of TAT-RasGAP_317-326_, meropenem and gentamicin. Error bars indicate standard deviations of triplicates. P values were calculated using two-way Anova in Graphpad Prism.

### 3.4 Combinations of three drugs can further improve bactericidal activity compared to combinations of two

To further decrease the concentrations of drugs needed to efficiently kill bacteria, we aimed at determining whether addition of a third compound would increase the death rate. In the case of *E. coli*, it was earlier demonstrated that addition of the Calcium chelator EDTA could increase the efficiency of TAT-RasGAP_317-326_ [20]. We thus combined TAT-RasGAP_317-326_, polymyxin B, and EDTA, and could observe that this combination was able to kill 99.9% of bacteria at concentrations that have no visible effect individually and limited activity in two-drug combinations (Fig. 3b). A similar effect was observed for *A. baumannii*, upon combination of TAT-RasGAP_317-326_ with both meropenem and gentamicin (Fig. 3c).

### 3.5 Combinations of TAT-RasGAP_317-326_ with gentamicin or meropenem cause robust additive growth inhibition on *A. baumannii* clinical isolates

Since combinations of TAT-RasGAP_317-326_ with gentamicin and/or meropenem had promising efficiency on the *A. baumannii* ATCC 19606 strains, we decided to test these combinations on a selection of clinical strains showing different levels of sensitivity towards meropenem and gentamicin. Three strains (17 0129 1317, 17 0112 2944 and 15 0815 0832) were selected from a former study in which they were shown to be sensitive to TAT-RasGAP_317-326_, four additional clinical isolates Ab31, Ab33, Ab44 and Ab73 were selected from a collection of clinical isolates originally described in [26] (Supplementary Table S1). Despite their diverse resistance profiles, all tested strains showed similar levels of sensitivity towards the antimicrobial peptides polymyxin B and TAT-RasGAP_317-326_ (Supplementary Table S1).

Regarding resistance to gentamicin, all isolates were resistant to gentamicin, EUCAST breakpoint of gentamicin being 4 μg/ml. However, resistance level was highly variable, from 8 to >256 μg/ml. We wondered whether combination of TAT-RasGAP_317-326_ with gentamicin could reverse resistance of these strains towards gentamicin. We could not observe a strong effect of this combination on the MIC of gentamicin in these clinical strains, using checkerboard assay (Table 2). Moreover, synergism between TAT-RasGAP_317-326_ and gentamicin could be detected only in one strain having an intermediate resistance to gentamicin similar to the ATCC19606 strain (Ab44, Table 2).

**Table 2:** Combination of TAT-RasGAP_317-326_ with gentamicin is additive on a majority of selected *A. baumannii* clinical isolates. Checkerboards with combinations of TAT-RasGAP_317-326_ and gentamicin were performed in triplicate on the indicated strains and MIC of the individual antimicrobial agents alone and in combination are presented. Cells containing two values indicate that all three replicates gave a value that falls within the indicated range.

| Strain ID | MIC Alone (µg/mL) |  | MIC Combination (µg/mL) |  | FICI |  |  |
| --- | --- | --- | --- | --- | --- | --- | --- |
|  | TAT-RasGAP<sub>317-326</sub> | Gentamicin | TAT-RasGAP<sub>317-326</sub> | Gentamicin |  |  |  |
| 15 0815 0832 | 8-32 | ≥ 256 | 1-4 | 128-256 | 0.625 | 0.625 | ≈ 0.625 |
| 17 0112 2944 | 32-64 | ≥ 256 | 16-32 | 64-256 | 0.75 | ≈ 1 | 1.25 |
| 17 0129 1317 | 16-64 | ≥ 256 | 4-32 | 64-128 | 0.625 | 0.75 | ≈ 0.75 |
| Ab31 | 32 | 8-16 | 4-16 | 4 | 0.5 | 0.625 | 0.75 |
| Ab33 | 8-32 | >256 | 8-32 | >256 | n.d. | n.d. | n.d. |
| Ab44 | 16-32 | 32 | 4-8 | 8 | 0.5 | 0.5 | 0.5 |
| Ab73 | 16-32 | 8-16 | 2-8 | 4-8 | 0.625 | 0.625 | 0.75 |

All seven clinical isolates were then challenged with combinations of TAT-RasGAP_317-326_ and meropenem (Table 3). MICs of meropenem towards the clinical isolates were highly variable (from 0.0625 to >256 μg/ml). Interestingly, combination of meropenem with TAT-RasGAP317-326 allowed to decrease MIC of meropenem between 2 and 16 times, approaching concentrations that can be reached in vivo, maximal serum concentration being approximately of 30 μg/ml [27] (Table 3).

**Table 3:** Combination of TAT-RasGAP_317-326_ with meropenem is at the limit between synergism and additivity on a majority of selected *A. baumannii* clinical isolates. Checkerboards with combinations of TAT-RasGAP_317-326_ and meropenem were performed in triplicate on the indicated strains and MICs of the individual antimicrobial agents alone and in combination are presented. Cells containing two values indicate that all three replicates gave a value that falls within the indicated range.

| Strain ID | MIC Alone (µg/mL) |  | MIC Combination (µg/mL) |  | FICI |  |  |
| --- | --- | --- | --- | --- | --- | --- | --- |
|  | TAT-RasGAP<sub>317-326</sub> | Meropenem | TAT-RasGAP<sub>317-326</sub> | Meropenem |  |  |  |
| 15 0815 0832 | 16-32 | 32-128 | 2-8 | 16-32 | 0.5 | 0.5 | 0.625 |
| 17 0112 2944 | 16-32 | ≥ 256 | 4-16 | 16 | 0.3125 | ≈ 0.625 | ≈ 0.625 |
| 17 0129 1317 | 32 | 4-8 | 8-16 | 0.125-1 | 0.5 | 0.5156 | 0.625 |
| Ab31 | 16-32 | 0.125-0.25 | 2-16 | 0.0313-0.125 | 0.5625 | 0.75 | 2 |
| Ab33 | 32 | 32-128 | 8 | 8-16 | 0.375 | 0.375 | 0.5 |
| Ab44 | 32 | 64-128 | 8 | 16-32 | 0.5 | 0.5 | 0.6125 |
| Ab73 | 8-16 | 0.0625 | 4-8 | 0.0078-0.0313 | 0.625 | 0.75 | 1 |

### 3.6 Combinations between TAT-RasGAP_317-326_ and gentamicin or meropenem have improved antibiofilm activity compared to individual drugs

Synergistic antibiofilm activity between antimicrobial peptides and antibiotics is described in the literature for some antimicrobial peptides such as colistin and melittin [28-30]. We thus investigated if combinations of TAT-RasGAP_317-326_ with classical antibiotics could inhibit biofilm formation and/or eradicate established biofilms. Firstly, we measured the individual effect of TAT-RasGAP_317-326_, gentamicin, and meropenem on biofilm formation and eradication of the *A. baumannii* ATCC 19606 strain using a protocol allowing *in vitro* biofilm formation, as described earlier [19] (Supplementary Fig. S6).

We then performed an adapted checkerboard assay to investigate the effect of combinations of TAT-RasGAP_317-326_ with gentamicin or meropenem. We measured the inhibition of biofilm formation (BFI, adding the drugs and bacteriasimultaneously) or the eradicationof established biofilm (BET, adding the drugs 24h after the induction of biofilm formation). Viability of bacteria composing the biofilm was estimated by measuring metabolic activity using resazurin (Res) and biofilm biomass was quantified by crystal violet (CV) staining. Combination of TAT-RasGAP_317-326_ and gentamicin showed an additive effect on bacterial viability during biofilm formation (FICI between ~ 0.3125 and ~ 0.625, Table 4 and Supplementary Fig. S7). However, gentamicin addition did not increase the effect of TAT-RasGAP_317-326_ on biomass accumulation (FICI ~ 1.5, Table 4 and Supplementary Fig. S7). In contrast, combination of TAT-RasGAP317-326 and gentamicin was not able to affect bacterial viability and biomass of established biofilms, even at the highest concentrations tested (Table 4 and Supplementary Fig. S8).

**Table 4:**
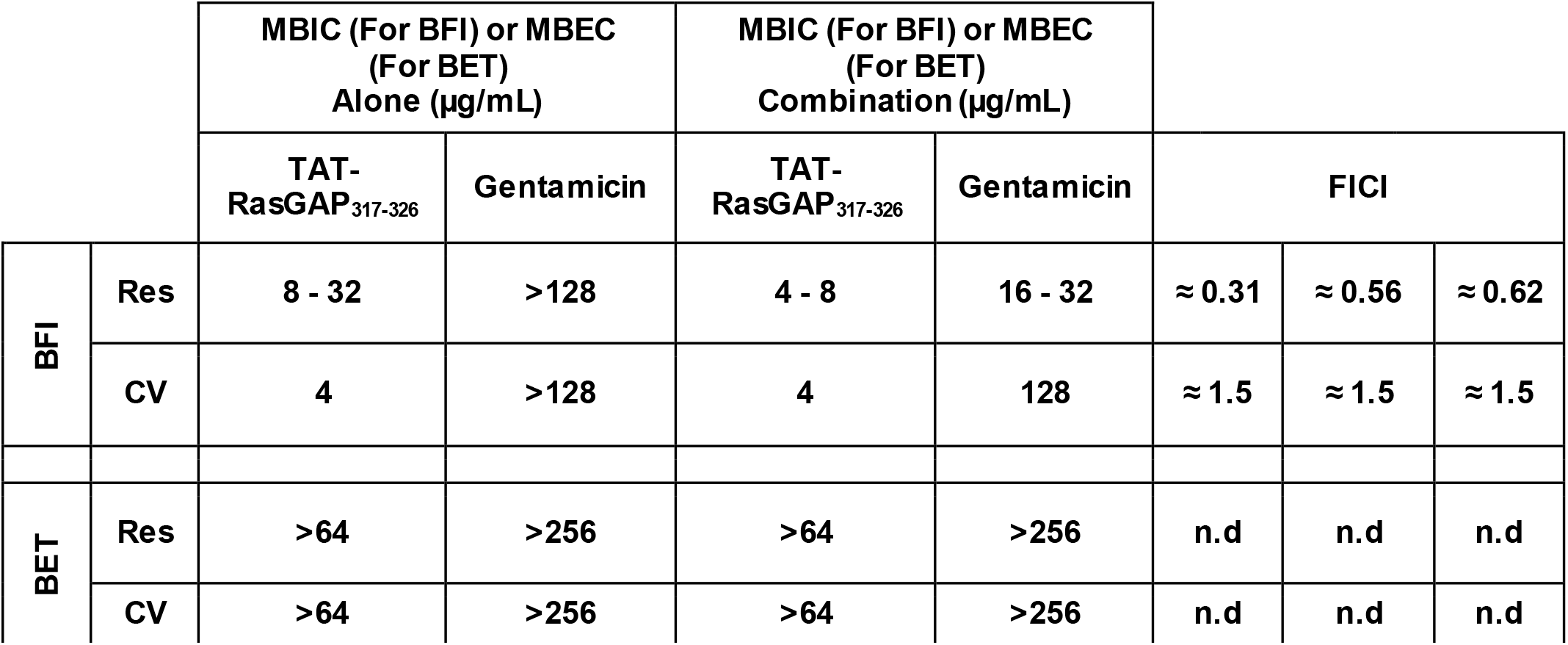
Combination of TAT-RasGAP_317-326_ with gentamicin has a strongly additive effect on viability of *A. baumannii* during biofilm formation, but is less efficient in inhibiting biofilm biomass formation and has no increased activity on established biofilms than individual antimicrobial agents. Checkerboard assays were performed in triplicate on forming biofilms (by adding the antimicrobial agents at the same time as bacterial seeding, BFI) and on biofilms preformed for 24 hours beforetreatment (BET). Bacterial viability was measured using resazurin (Res) and biofilm biomass was evaluated by crystal violet staining (CV). Complete checkerboard results are shown in Supplementary Figures S7 and S8. Minimal biofilm inhibitory concentration (MBIC), respectively Minimal biofilm eradication concentration (MBEC) are shown for individual antimicrobial agents and for combination. MBIC and MBEC were calculated independently for viability readout (Res) and for biomass readout (CV). Cells containing two values indicate that all three replicates gave a value that falls within the indicated range. n.d.: not determined.

|  |  | MBIC (For BFI) or MBEC (For BET) Alone (µg/mL) |  | MBIC (For BFI) or MBEC (For BET) Combination (µg/mL) |  |  |  |  |
| --- | --- | --- | --- | --- | --- | --- | --- | --- |
|  |  | TAT-RasGAP<sub>317-326</sub> | Gentamicin | TAT-RasGAP<sub>317-326</sub> | Gentamicin | FICI |  |  |
| BFI | Res | 8 - 32 | >128 | 4 - 8 | 16 - 32 | ≈ 0.31 | ≈ 0.56 | ≈ 0.62 |
|  | CV | 4 | >128 | 4 | 128 | ≈ 1.5 | ≈ 1.5 | ≈ 1.5 |
| BET | Res | >64 | >256 | >64 | >256 | n.d | n.d | n.d |
|  | CV | >64 | >256 | >64 | >256 | n.d | n.d | n.d |

Conversely, combination of TAT-RasGAP_317-326_ with meropenem synergistically affected bacteria viability in forming biofilms (FICI between 0.2531 and 0.375, Table 5 and Supplementary Fig. S9) and efficiently hindered the accumulation of biofilm biomass (FICI between ~ 0.1875 and ~ 0.5156, Table 5 and Supplementary Fig. S9). A synergistic effect on bacterial viability of the combination was also observed in established biofilms (FICI between ~ 0.132 and ~ 0.2813, Table 5 and Supplementary Fig. S10), but no decrease of biomass was observed on established biofilms, indicating that the combination was not able to disrupt established biofilm architecture (Table 5 and Supplementary Fig. S10).

**Table 5.** Combination of TAT-RasGAP_317-326_ with meropenem has a synergistic effect on viability of *A. baumannii* during biofilm formation and in preformed biofilms, a strong additive inhibitory effect on forming biofilm biomass and has no increased activity on biomass of established biofilms than individual antimicrobial agents. Checkerboard assays were performed in triplicate on forming biofilms (by adding the antimicrobial agents at the same time as bacterial seeding, BFI) and on biofilms preformed for 24 hours before treatment (BET). Bacterial viability was measured using resazurin (Res) and biofilm biomass was evaluated by crystal violet staining (CV). Complete checkerboard results are shown in Supplementary Figures S9 and S10. Minimal biofilm inhibitory concentration (MBIC), respectively Minimal biofilm eradication concentration (MBEC) are shown for individual antimicrobial agents and for combination. MBIC and MBEC were calculated independently for viability readout (Res) and for biomass readout (CV). Cells containing two values indicate that all three replicates gave a value that falls within the indicated range. n.d.: not determined.

|  |  | MBIC (For BFI) or MBEC<br>(For BET)<br>Alone (µg/mL) |  | MBIC (For BFI) or MBEC<br>(For BET)<br>Combination (µg/mL) |  |  |  |  |
| --- | --- | --- | --- | --- | --- | --- | --- | --- |
|  |  | TAT-RasGAP<sub>317-326</sub> | Meropenem | TAT-RasGAP<sub>317-326</sub> | Meropenem | FICI |  |  |
| BFI | Res | 16 | 32 | 4 | 1-4 | 0.2531 | 0.3125 | 0.375 |
|  | CV | 4-8 | >128 | 1-2 | 2-16 | ≈ 0.19 | ≈ 0.31 | ≈ 0.52 |
| BET | Res | 64->128 | 128->256 | 16-32 | 4-16 | ≈ 0.13 | ≈ 0.28 | 0.2813 |
|  | CV | >256 | >256 | >256 | >256 | n.d. | n.d. | n.d. |

## 4. Discussion

TAT-RasGAP_317-326_, like many AMPs, is a potent antimicrobial agent with a broad spectrum of activity [18]. However, its pharmacokinetic properties and its rapid excretion by the organism, linked to the high concentrations required for efficient antimicrobial activity, hinder this peptide’s clinical application. Alternative strategies are thus needed to improve TAT-RasGAP_317-326_ efficiency. One possible strategy is the development of combinatory treatments that would result in synergistic effects, which implies that the total effect is greater than the expected additive effect of individual drugs thus allowing an efficient antimicrobial activity with relatively low concentrations of each antimicrobial agent. Here, we identified synergistic interactions between TAT-RasGAP_317-326_ and common antimicrobial agents for all three bacterial species tested. Interestingly, the effect of combinations between antimicrobial agents was different for each bacterial species we tested. Namely, only polymyxin B had a strong interaction with TAT-RasGAP_317-326_, substantially reducing the concentration of TAT-RasGAP_317-326_ needed to inhibit *E. coli* visible growth. Similarly, against *P. aeruginosa*, the antimicrobial activity of TAT-RasGAP_317-326_ was increased in presence of aztreonam and polymyxin B. By contrast, several antimicrobial agents potently increased the activity of TAT-RasGAP_317-326_ on *A. baumannii*. However, these observations are based on a preliminary screening and should be confirmed with further investigations in the future.

Synergistic interaction between three different antimicrobial agents, polymyxin B, aztreonam, and gentamicin, and TAT-RasGAP_317-326_ on *E. coli, P. aeruginosa*, and *A. baumannii*, respectively, were confirmed. The relatively high FICI values (between 0.375 and 0.5) that were obtained indicate that the effect of these combinations may be further improved. This is consistent with the suboptimal bactericidal effect of combinations we measured (Fig. 2). We could increase the bactericidal effect of combinations by adding either EDTA, a Calcium chelator against *E. coli* or by adding a third antimicrobial agent towards *A. baumannii*. Such multicomponent combinations need to be further studied to determine the optimal combinations that may induce the best bactericidal effect with the lowest concentrations of antimicrobial agents.

The clinical relevance of combinations tested here has still to be confirmed. Indeed, concentrations of the individual components need to be reachable *in vivo*. Interestingly, concentrations of aztreonam and polymyxin B used in combinations against *P. aeruginosa* and *E. coli*, respectively, are well below the serum maximal concentrations described in the literature (Supplementary Table S2) [31],[32]. This is especially interesting for clinical strains of *A. baumannii* that show a low resistance to gentamicin (8-16 μg/ml). Combination with TAT-RasGAP_317-326_ allows to increase their sensitivity to gentamicin to concentrations that are achievable in the serum (Table 2) [33]. This is also true for meropenem, for which the maximal serum concentration is approximately of 30 μg/ml [27]. The main limitation is the limited bioavailability of TAT-RasGAP_317-326_, which shows a maximal serum concentration of approximately 2.8 μg/ml in a mouse model [21]. This limits potential systemic applications of combinatory therapies using TAT-RasGAP_317-326_. However, topical administration may allow to reach the required concentrations. This is also the case for EDTA, for which concentrations up to 0.1 % are used for topical treatment. In contrast, intravenous administration of EDTA is used to treat heavy metal intoxications, but can lead to important side effects, limiting the possibility of treatment of systemic bacterial infections with combinations containing high concentrations of EDTA.

In the environment or during infection, bacteria are often part of biofilms. AMPs are cited as especially potent antibiofilm molecules. TAT-RasGAP_317-326_ can inhibit the formation of *A. baumannii* biofilm *in vitro*, but is much less efficient on established biofilms [19]. No improvement was observed by combination of TAT-RasGAP_317-326_ with gentamicin (Table 4). However, combination of this peptide with meropenem decreased the viability of bacteria in established biofilms, but not the biofilm biomass (Table 5). This is reminiscent of the difficulties to develop agents that efficiently eradicate established biofilms [34].

In this article, we described synergistic combinations of TAT-RasGAP_317-326_ with commonly used antimicrobial agents on three Gram-negative bacteria. We show that combinations can efficiently inhibit growth of *A. baumannii* clinical isolates and can affect bacterial viability in established *A. baumannii* biofilms. Antimicrobial effect of these combinations needs to now be tested *in vivo* to confirm their potential and hopefully lead to the development of treatment strategies that may allow to treat infections caused by MDR bacteria. Finally, combinatory treatments are believed to reduce the risk of resistance emergence. However, broad range resistance to AMPs is still possible [35]. This needs to be investigated in more detail regarding combinations with TAT-RasGAP_317-326_.

## Abbreviations

AMPs: antimicrobial peptides
MDR: multidrug resistant
MIC: Minimal inhibitory concentration
CFU: Colony forming units
FICI: Fractional inhibitory concentration index
MBIC: Minimal biofilm inhibitory concentration
MBEC: Minimal biofilm eradication concentration
IC50: Inhibitory concentration 50%

## Acknowledgements

We thank Prof. Gilbert Greub and the Institute of Microbiology for providing laboratory space and equipment, as well as clinical isolates. We also thank Dr. Gregory Resch and Prof. Leo Eberl for kindly sharing some strains, Sébastien Aeby and Hugo de Jesus for technical support. We finally thank Simone Hargraves for critical reading of this manuscript. This work was partially financed by an interdisciplinary grant from the Faculty of Biology and Medicine of the University of Lausanne.

## Author contributions

NJ designed the project, GV and NJ planned and performed the experiments. NJ and GV analysed the data, prepared the figures and wrote the manuscript.

## Data availability statement

All data produced for this study are included in the Figures, Tables and Supplementary data. Raw data can be available upon request to the corresponding author.

## Additional information

The authors declare no competing interests.

