## Supplementary data for "*In vitro* synergistic action of TAT-RasGAP_317-326_ peptide with antibiotics against Gram-negative pathogens"

**Table S1: Minimal inhibitory concentrations (MIC) and concentrations causing a growth inhibition of 50% (IC50) of the antimicrobial agents used in this study on laboratory and clinical strains of *Escherichia coli*, *Pseudomonas aeruginosa* and *Acinetobacter baumannii* (in µg/mL).** Measurements were performed in duplicate. Two values of MIC are shown when both measurements resulted in different values. $IC_{50}$ values were calculated using Graphpad Prism on the average of the two replicates.

| Strain name | Reference | Azithromycin | Aztreonam | Ceftriaxone | Ciprofloxacin | Erythromycin | Gentamicin | Meropenem | Tetracycline | Polymyxin B | TAT-RasGAP$_{317-326}$ |
|---|---|---|---|---|---|---|---|---|---|---|---|
| *E. coli* MG1655 (ATCC 47076) | ATCC | 32 | 0.0625 | 0.0625 | 0.03125 | >256 | 32-64 | 0.0625 | 2 | 2 | 16-32 |
| *E. coli* ATCC 25922 | ATCC | 16-32 | 0.0625-0.125 | 0.03125 | 0.0078125 | 256 | 16-32 | 0.03125 | 1 | 4 | 32-64 |
| *P. aeruginosa* PA14 | Rahme et al., 1995 | >256 | 8 | 16 | 0.125 | >256 | 2 | 0.5 | 8 | 2 | 64-128 |
| *P. aeruginosa* PA01 | Stover et al., 2000 | >256 | 8-16 | 32-64 | 0.125 | >256 | 02.avr | 0.5 | 16 | 4-8 | 128 |
| *A. baumannii* ATCC 19606 | ATCC | 32-64 | 16-32 | 32-64 | 1 | 256 | 64 | 1-2 | 2 | 2 | 16 |
| *A. baumannii* 15 0815 0832 | Heulot et al., 2017 | 32 | 32-64 | >256 | 64 | 64 | >256 | 64 | >256 | 2 | 8-16 |
| *A. baumannii* 17 0112 2944 | Heulot et al., 2017 | >256 | 128 | >256 | 64 | >256 | >256 | >256 | 1 | 2 | 16 |
| *A. baumannii* 17 0129 1317 | Heulot et al., 2017 | 64-128 | 16 | >256 | 256 | 32 | >256 | 16 | >256 | 4 | 8-16 |
| *A. baumannii* Ab31 | Leshkasheli et al., 2019 | 1 | 32 | 16 | 8 | 16 | 16 | 0.25 | 1 | 4 | 16 |
| *A. baumannii* Ab33 | Leshkasheli et al., 2019 | 2 | 32 | >256 | >256 | >256 | >256 | 64 | >256 | 2 | 16 |
| *A. baumannii* Ab44 | Leshkasheli et al., 2019 | >256 | 128 | >256 | >256 | >256 | 16-32 | 256 | 2 | 4 | 16 |
| *A. baumannii* Ab73 | Leshkasheli et al., 2019 | 1 | 64 | 8 | 8 | 8 | 8 | 0.125 | 1 | 2 | 16 |

**Table S2: List of antimicrobial agents used in this study with their family, mode of action, EUCAST breakpoints and approximative serum maximal concentrations.** S: Sensitive, R: Resistant N. d.: not determined. (1) Breakpoints represent MICs (mg/L) from EUCAST breakpoint tables version 12.0. (2) Vinks *et al.*, *Antimicrob Agents Chemother*, 2007. (3) Kovacevic *et al.*, *J Basic Clin Pharm*, 2016. (4) Moon *et al.*, *Clin Infect Dis*, 1997. (5) Not available from EUCAST, values from USCAST, Pogue *et al.*, *Antimicrob Agents Chemother*, 2020. (6) Sandri *et al.*, *Clin Infec Dis*, 2013.

| Drug | Family | Mode of Action | *E. coli* breakpoints[1] | | *P. aeruginosa* breakpoints[1] | | *A. baumannii* breakpoints[1] | | Non-species related breakpoints[1] | | Approximative serum maximal concentration |
|---|---|---|---|---|---|---|---|---|---|---|---|
| | | | S ≤ | R > | S ≤ | R > | S ≤ | R > | S ≤ | R > | |
| **AZITHROMYCIN** | MACROLIDES | Translation inhibition | n.d | n.d. | n.d | n.d. | n.d | n.d. | n.d | n.d. | |
| **AZTREONAM** | MONOBACTAMS | Inhibition of cell wall synthesis | 1 | 4 | 0.001 | 16 | n.d | n.d. | 4 | 8 | >100 (2) |
| **CEFTRIAXONE** | CEPHALOSPORINS | Inhibition of cell wall synthesis | 1 | 2 | n.d | n.d. | n.d | n.d. | 1 | 2 | |
| **CIPROFLOXACIN** | FLUOROQUINOLONES | DNA-gyrase inhibitors | 0.25 | 0.5 | 0.001 | 0.5 | 0.001 | 1 | 0.25 | 0.5 | |
| **ERYTHROMYCIN** | MACROLIDES | Translation inhibition | n.d | n.d. | n.d | n.d. | n.d | n.d. | n.d | n.d. | |
| **GENTAMICIN** | AMINOGLYCOSIDES | Translation inhibition | 2 | 2 | n.d | n.d. | 4 | 4 | 0.5 | 0.5 | 5 to 10 (3) |
| **MEROPENEM** | CARBAPENEMS | Inhibition of cell wall synthesis | 2 | 8 | 2 | 8 | 2 | 8 | 2 | 8 | 30 (4) |
| **TETRACYCLINE** | TETRACYCLINES | Translation inhibition | n.d | n.d. | n.d | n.d. | n.d | n.d. | n.d | n.d. | |
| **POLYMYXIN B** | CYCLIC POLYPEPTIDES | Membrane disruption | n.d | n.d. | n.d | n.d. | n.d | n.d. | 2 (5) | 4 (5) | 5 to 10 (6) |

**Figure S1: Concentrations of antimicrobials selected for initial screening of combinations cause a partial growth defect of the target bacteria.** $OD_{590}$ was measured after 16 hours of incubation of *E. coli* MG1655 **(a)**, *P. aeruginosa* PA14 **(b)** and *A. baumannii* ATCC 19606 **(c)** in presence of the indicated concentrations of antimicrobial agents and is represented as percentage of growth to an untreated control.

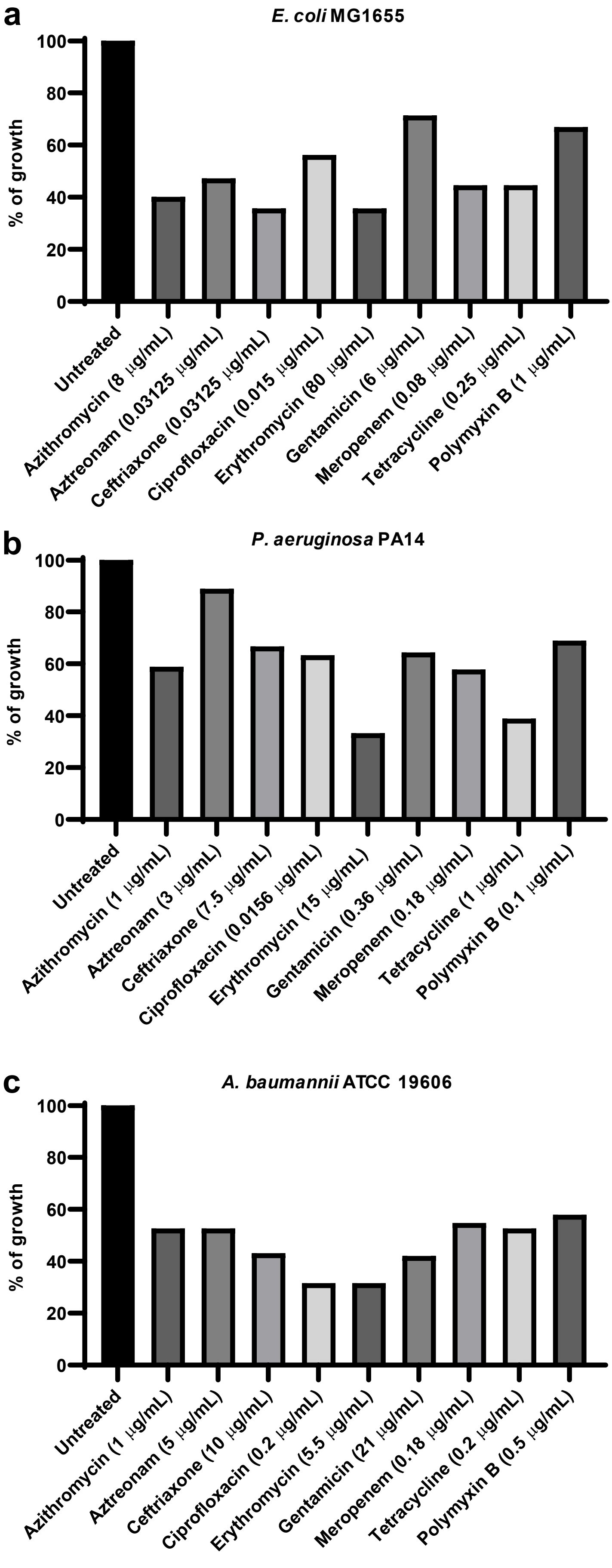

**Figure S2: Checkerboard assays with combinations of TAT-RasGAP$_{317-326}$ and polymyxin B on *E. coli* MG1655 (a) and ATCC15922 (b).** The indicated strains were grown for 16 hours at 37°C in presence of serial dilutions of the drugs, as indicated. OD$_{590}$ was measured and is shown as color gradient. Results of three independent experiments are shown for each strain.

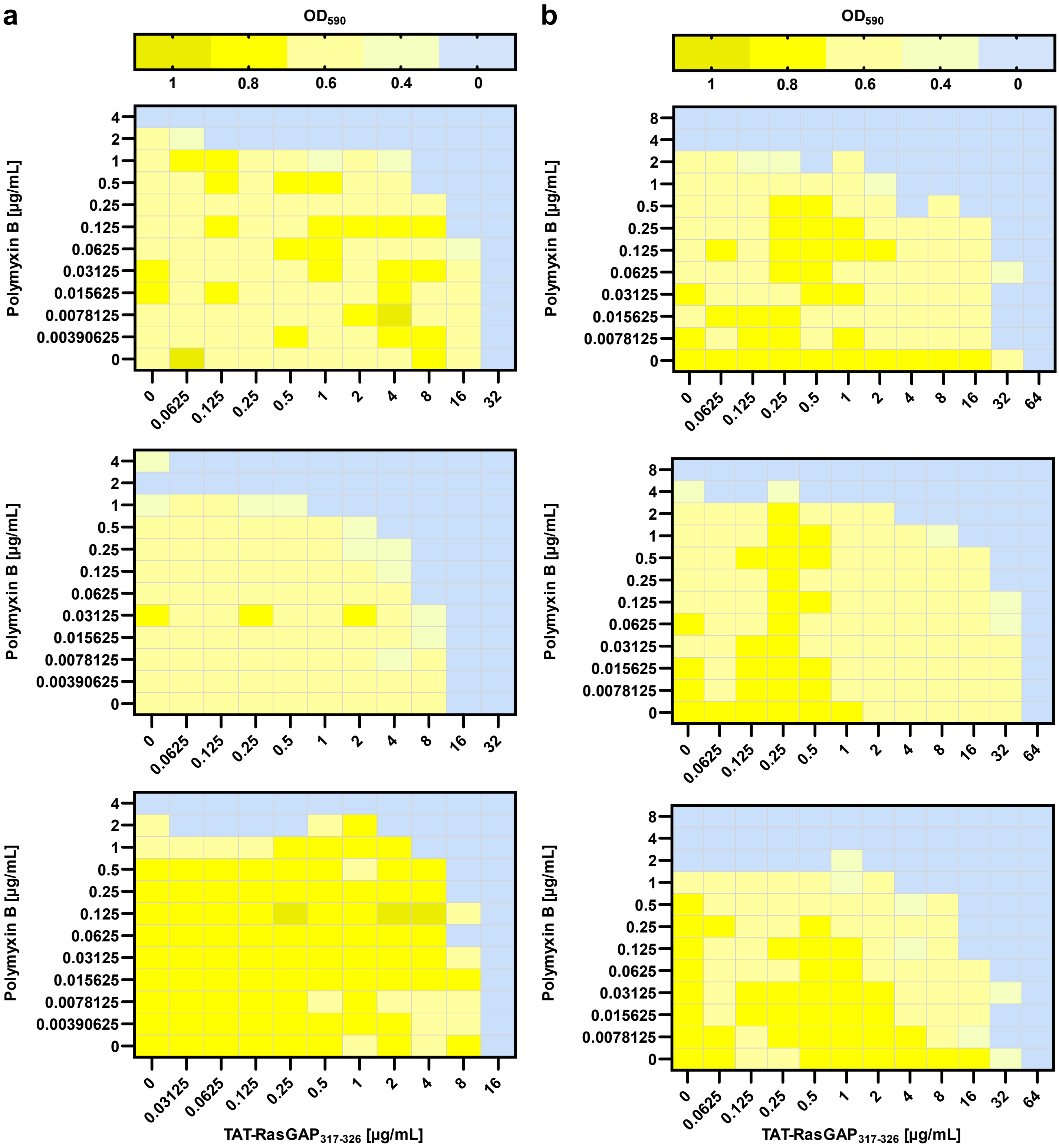

**Figure S3: Checkerboard assays with combinations of TAT-RasGAP$_{317-326}$ and aztreonam on *P. aeruginosa* PA14 (a) and PA01 (b).** The indicated strains were grown for 16 hours at 37°C in presence of serial dilutions of the drugs, as indicated. $OD_{590}$ was measured and is shown as color gradient. Results of three independent experiments are shown for each strain.

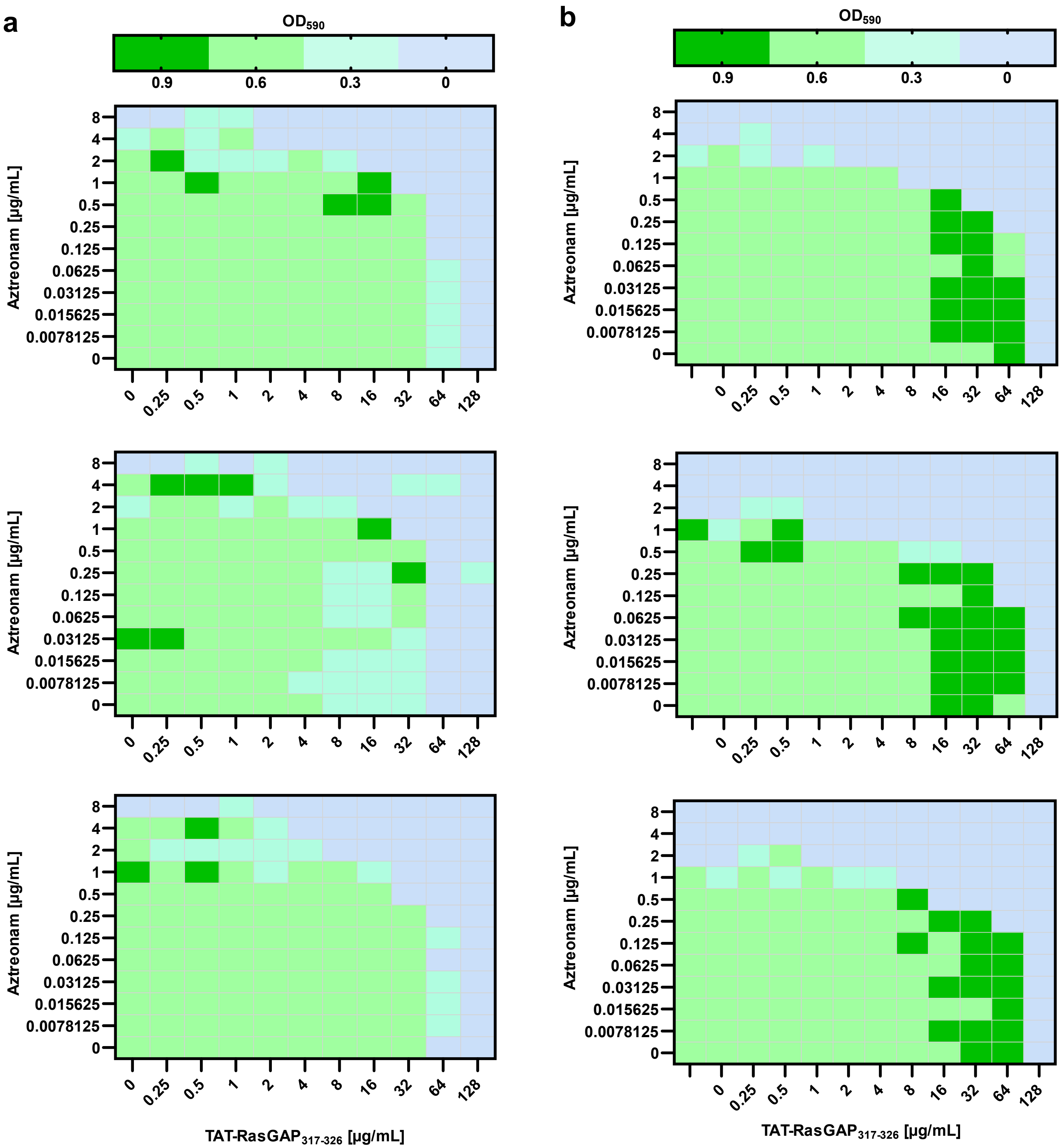

**Figure S4: Checkerboard assays with combinations of TAT-RasGAP$_{317-326}$ and gentamicin (a) or meropenem (b) on *A. baumannii* ATCC 19606.** The indicated strains were grown for 16 hours at 37°C in presence of serial dilutions of the drugs, as indicated. $OD_{590}$ was measured and is shown as color gradient. Results of three independent experiments are shown for each condition.

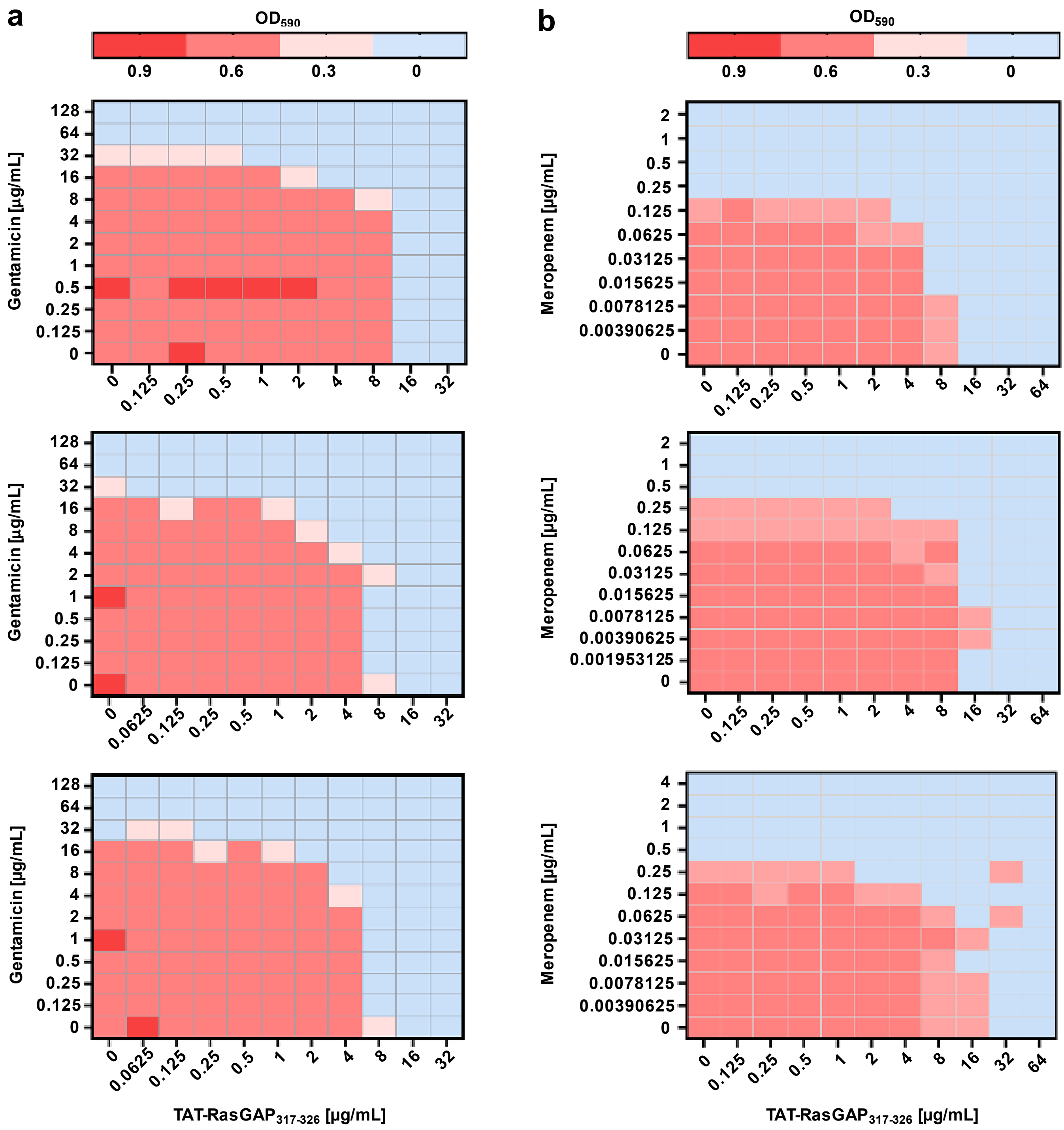

**Figure S5: Foldchange of CFU counts with increasing concentrations of individual antimicrobial agents.** Overnight culture of *E. coli* MG1655, *P. aeruginosa* PA14 or *A. baumannii* ATCC19606 were diluted to 0.1 $OD_{600}$ and grown for one hour at 37°C before the addition of the indicated concentrations of antimicrobial agents. Incubation was then continued and samples were taken at 0h and 2h after addition of the drugs. Serial dilutions of these samples were plated on LB agar plates and number of Colony Forming Units per ml (CFU/mL) were quantified. Fold change between CFU/mL of each time point and the initial value (time 0) was calculated and is shown as $Log_{10}$ values. Horizontal dotted line indicates the threshold below which more than 99.9% of bacteria are dead. Red numbers highlight, when available, the minimal concentrations allowing at least 99.9% of death ($MBC_{99.9}$). Error bars indicate standard deviation of triplicates.

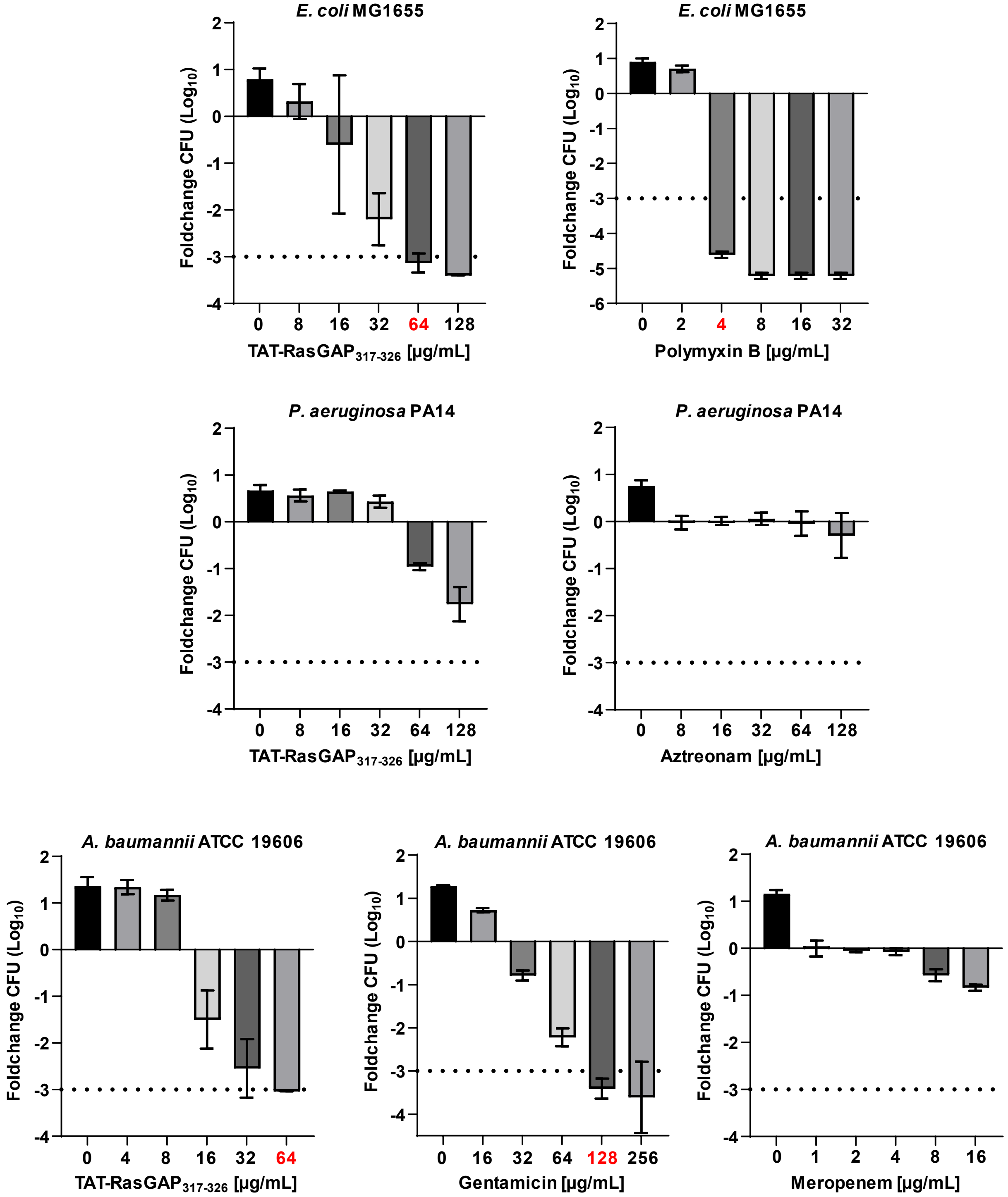

**Figure S6: Effect of TAT-RasGAP$_{317-326}$ (a), gentamicin (b) and meropenem (c) on biofilm formation and on mature biofilms of *A. baumannii* ATCC 19606.** Biofilms were prepared as described in Fig. 4 with serial dilutions of the indicated antimicrobial agents added at the time of seeding (Biofilm inhibition) or after 24h (Biofilm eradication). Viability of bacteria composing the biofilm was measured 24h after addition of the antimicrobial agents using resazurin (Res) and biofilm biomass was estimated using crystal violet staining (CV).

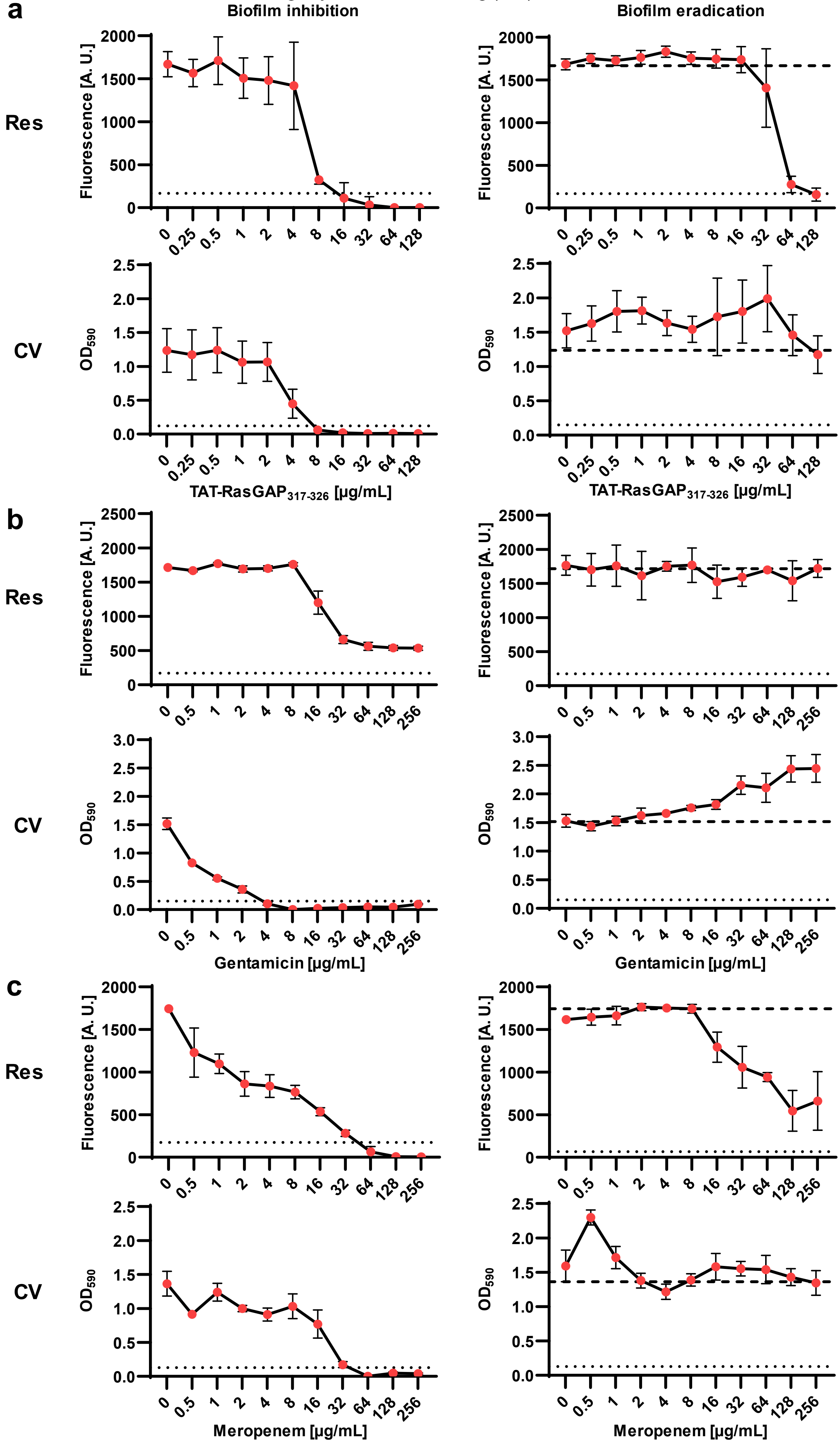

**Figure S7: Effect of combination between TAT-RasGAP$_{317-326}$ and gentamicin on biofilm formation.** Checkerboard assays were performed on forming biofilm by adding the antimicrobial agents at the same time as bacterial seeding. Bacterial viability was measured using resazurin and biofilm biomass was evaluated by crystal violet staining 24 hours after drug addition. Results of three independent experiments are shown. Fluorescence and $OD_{590}$ were measured, respectively, and are shown as color gradients.

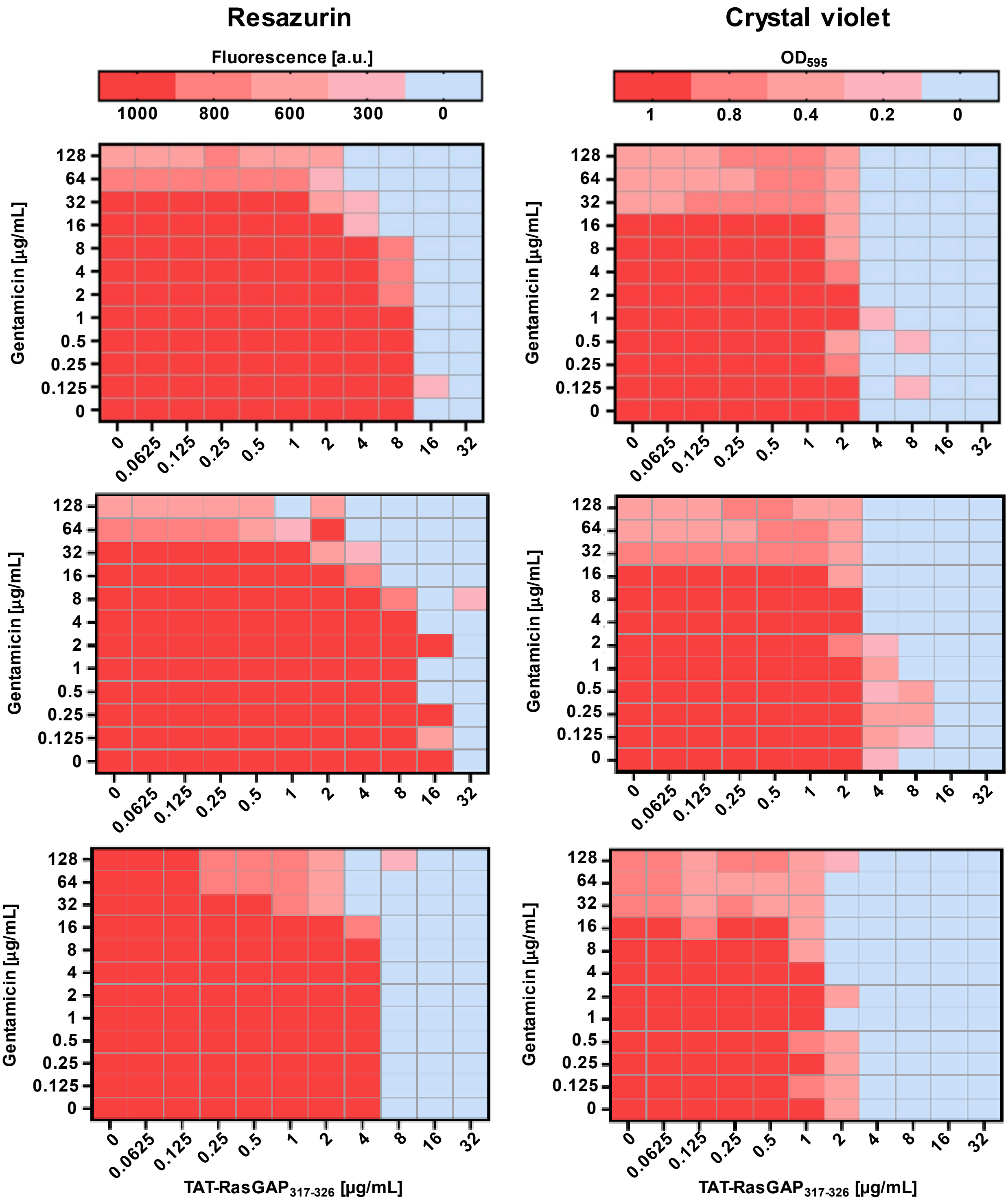

**Figure S8: Effect of combination between TAT-RasGAP$_{317-326}$ and gentamicin on mature biofilms.** Checkerboard assays were performed on mature biofilms by adding the antimicrobial agents 24h after the initiation of biofilm formation. Bacterial viability was measured using resazurin and biofilm biomass was evaluated by crystal violet staining 24 hours after drug addition. Results of three independent experiments are shown. Fluorescence and $OD_{590}$ were measured, respectively, and are shown as color gradients.

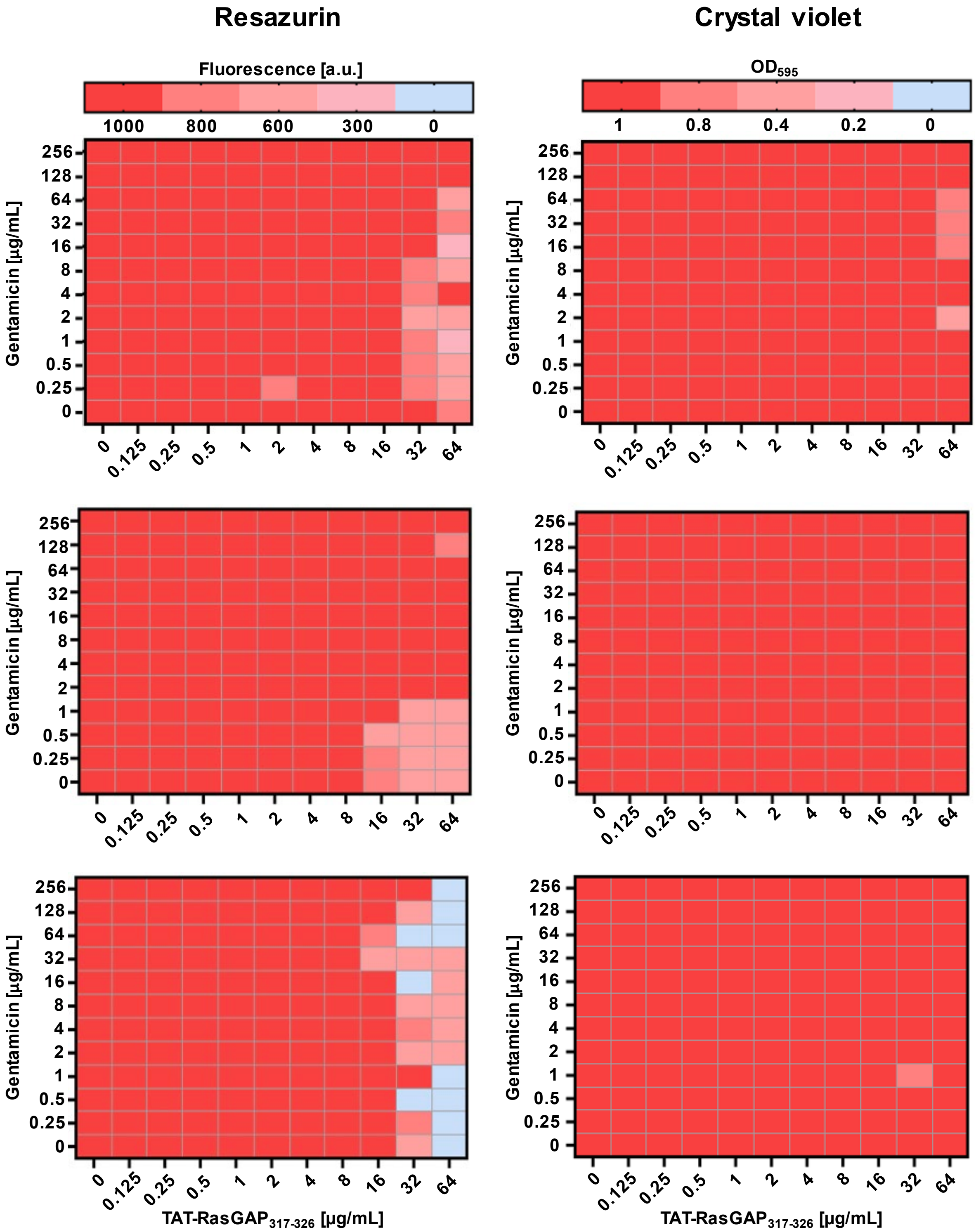

**Figure S9: Effect of combination between TAT-RasGAP$_{317-326}$ and meropenem on biofilm formation.** Checkerboard assays were performed on forming biofilms by adding the antimicrobial agents at the same time as bacterial seeding. Bacterial viability was measured using resazurin and biofilm biomass was evaluated by crystal violet staining 24 hours after drug addition. Results of three independent experiments are shown. Fluorescence and OD$_{590}$ were measured, respectively, and are shown as color gradients.

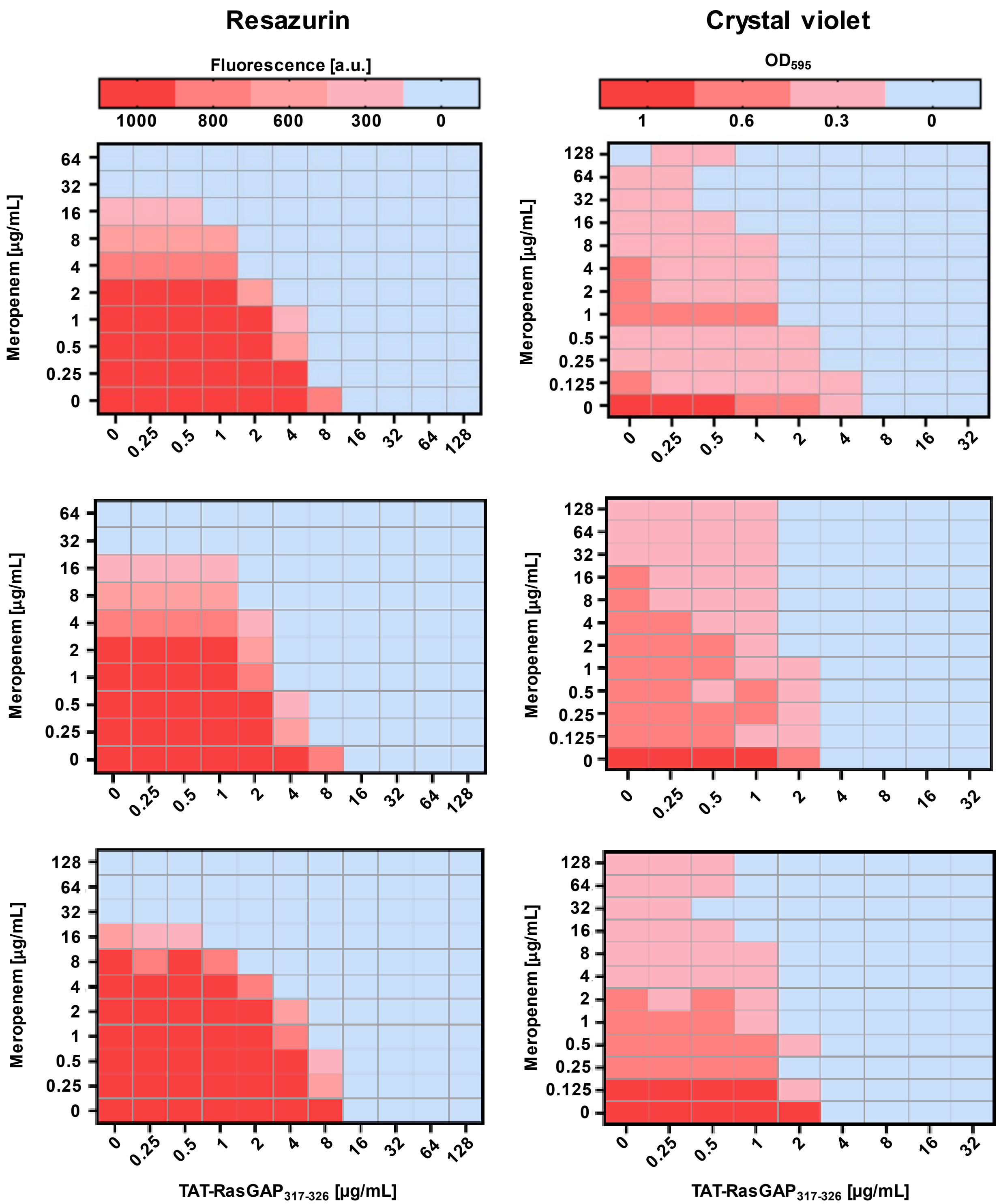

**Figure S10: Effect of combination between TAT-RasGAP$_{317-326}$ and meropenem on mature biofilms.** Checkerboard assays were performed on mature biofilms by adding the antimicrobial agents 24h after the initiation of biofilm formation. Bacterial viability was measured using resazurin and biofilm biomass was evaluated by crystal violet staining 24 hours after drug addition. Results of three independent experiments are shown. Fluorescence and OD$_{590}$ were measured, respectively, and are shown as color gradients.

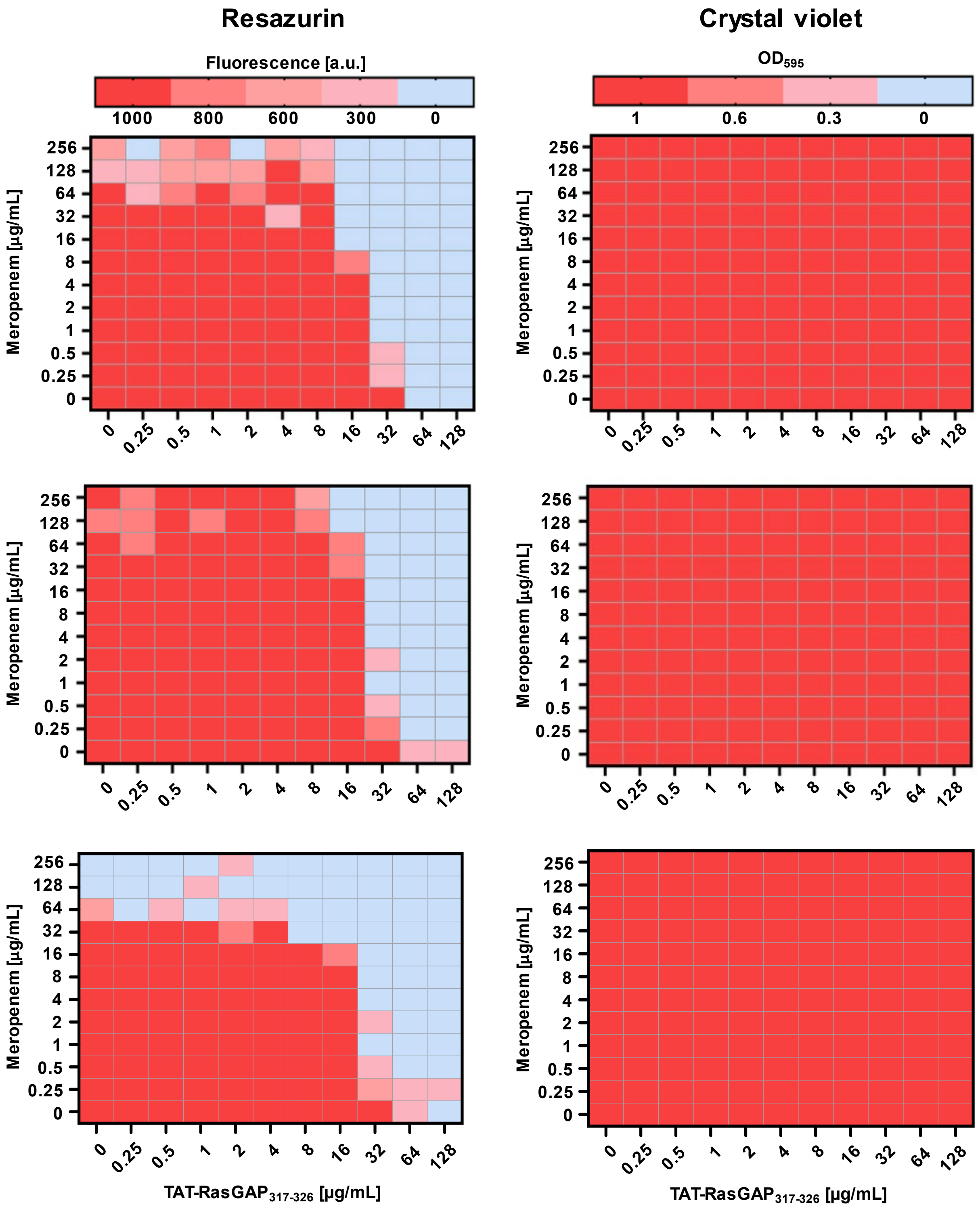
